# Ontogeny of escape response and body shape in Threespine Stickleback (*Gasterosteus aculeatus* L.)

**DOI:** 10.1101/2025.07.09.663780

**Authors:** Aspen M. Kozak, Michael Chung, Sean M. Rogers, Kelsey N. Lucas, Heather A. Jamniczky

**Author notes:** Corresponding author: Heather Jamniczky Cumming School of Medicine University of Calgary, 3330 Hospital Dr NW Calgary AB T2N 4N1. Contributed equally (Kelsey N. Lucas, Heather A. Jamniczky).

## Abstract

Escape responses in fishes provide insight into accelerative motions and behavioral response times of these animals, linking numerous fitness-related traits. We sought to connect escape response performance to genotype and phenotype across ontogenetic stages within a single population of Threespine Stickleback (*Gasterosteus aculeatus* L.), to determine if a gene of major effect (Ectodysplasin; *Eda*) on adaptive lateral armour plate phenotype and fitness conveys a performance advantage in brackish and saltwater environments. We predicted that possession of one or more low-plated alleles would result in better escape performance in brackish environments across ontogeny, and that *Eda* genotype would result in variation in shape phenotype that would be predictive of escape performance before the completion of lateral plate development. We assessed three kinematic variables derived from C-start escape responses in wild-caught, F1 juvenile, and F1 adults, and found that *Eda* genotypes did not directly impact maximum velocity or curvature coefficient, but linked phenotypes contributed to an increase in total distance traveled in F1 adults. We further showed that *Eda* genotype impacted growth, producing variation in size which in turn influenced escape performance variables across age groups, with a particular focus on total distance traveled. The influence of *Eda* genotype on shape was confounded by sexual dimorphism, indicating that further sex-specific studies on shape and performance are needed. The presence of at least one low-plated allele conferred a growth advantage, particularly in a freshwater rearing environment, consistent with known pleiotropic effects of *Eda*, including on skeletal development, that may be modulating anti-predator responses before plate phenotype is fully expressed. Altogether, more ontogenic studies are needed to ensure the influence of development on adaptive traits is more clearly understood.

## Introduction

Organismal structure and function are closely related and are in part reflective of the physical constraints of an organism’s environment. Performance traits will inevitably affect individual fitness and contribute to shaping the evolution of populations and ecosystems. Understanding how organisms adapt to rapidly changing environments, such as those impacted by anthropogenic influence like urbanization and shoreside construction, will therefore require extension of biomechanical analyses past the individual organism to link ecological performance on a population or community scale to environmental context (Higham et al., 2021; Higham et al., 2016). While unlikely to provide an accurate quantitative measure of fitness because they do not usually account for all factors related to reproductive success, performance measures can be useful in generating qualitative inferences about natural selection (Franklin et al., 2017) and provide an important link between morphology and fitness (Arnold, 1983) that allows us to explore the effects of mutations that underlie rapid adaptive change. Large-effect mutations that impact fitness have been proposed to be beneficial under certain environmental conditions (Orr, 2005), and the *Heliconius* butterfly is one such example in a contemporary context (Heliconius Genome, 2012). Studies documenting the fitness effects of large-effect mutations in wild populations across generations, however, remain rare (Schluter et al., 2021).

The threespine stickleback fish (*Gasterosteus aculeatus*; stickleback) provides an intriguing model to study the relationship between performance and adaptation in fishes, and the impact of large-effect mutations on evolvability and adaptive capacity in a contemporary example of parallel evolution. Stickleback have repeatedly colonized newly formed freshwater habitat across the Holarctic since the retreat of the Pleistocene glaciers (Bell & Foster, 1994) and continue to do so on a contemporary timescale (Kozak et al., 2024). Newly established freshwater populations repeatedly and consistently diverge from marine populations in a variety of morphological, behavioural, physiological, genetic, and life history traits. A striking feature of parallel evolution in stickleback is the well documented loss of lateral armour plates across the salt-freshwater transition (e.g. Colosimo et al., 2005; Schluter et al., 2004) which is driven by a large-effect mutation in *Ectodysplasin* (*Eda*). Complete plate morphs, with a continuous row of 30-36 lateral plates spanning the length of the body, represent the putative ancestral condition that dominates in marine environments. Low plate morphs, where nine or fewer anterior plates are retained, represent the derived condition that dominates in freshwater environments. Partial plate morphs, describing individuals with either a gap of at least two plates or an intermediate number of plates, are rare in both marine and freshwater environments (Barrett et al., 2008; Bell, 2001; Hagen & Gilbertson, 1972).

*Eda* behaves like a Mendelian factor (Cresko et al., 2004) and explains more than 70% of the variation in number of lateral plates (Colosimo et al., 2005). Homozygotes for the complete-plated allele develop into complete morphs, homozygotes for the low-plated allele develop into low morphs, and heterozygotes develop into partial morphs or complete morphs. The expression of lateral plates and associated spines is reduced by a cis-regulatory change in the low-plated *Eda* haplotype (O’Brown et al., 2015). The *Eda* receptor (*Edar*) has also recently been shown to impact plate number variation (Laurentino et al., 2022). Identical low-plated *Eda* haplotypes have been found in freshwater populations across the Holarctic (Colosimo et al., 2005; O’Brown et al., 2015), suggesting that the low-plated *Eda* haplotype exists as standing genetic variation in marine populations, and the low-plated *Eda* allele has been shown to persist at a frequency of ∼1% in marine populations (Barrett et al., 2008; Colosimo et al., 2005; Morris et al., 2018). *Eda* has additional, well-documented pleiotropic effects on body shape (Aguirre & Bell, 2012; Albert et al., 2008; Rogers et al., 2012), schooling behaviour (Greenwood et al., 2013) and lateral line neuromast distribution in stickleback (Mills et al., 2014), which emerge from its roles in the development of bone and neural tissue (Rodriguez-Ramirez et al., 2023).

There is ample evidence that the low-plated phenotype is adaptive in freshwater environments (Barrett, 2010). However, there remains a lack of consensus as to the precise nature of the environmental or intrinsic force(s) associated with selection on *Eda*. Hypotheses include physiological differences in salt tolerance resulting in selection of osmoregulatory genes near *Eda* (Heuts, 1947), increased cost of plate production due to calcium and/or phosphorous limitation on skeletal development (Archambeault et al., 2020; Giles, 1983), a trade-off between plate production and growth resulting in low morphs having a growth advantage in freshwater (Marchinko & Schluter, 2007) and correspondingly higher overwinter survival (Barrett et al., 2008), and temperature effects (Kanbe et al., 2023; Smith et al., 2021).

Changes in predation regime have also been hypothesized as a driving factor for differential selection on stickleback (Barrett, 2010; Leinonen et al., 2011; Reimchen, 1994), but the relationship between stickleback and their predators, and the resulting impact on selection, is complex (Reimchen & Bergstrom, 2023). The presence of a complete row of lateral plates reduces the extent of injuries during capture, particularly with toothed predators, and increases the chance of predator handling failure and stickleback escape from capture (Reimchen, 1992, 1994, 2000; Reimchen & Nosil, 2004). These findings support the hypothesis that complete morphs are favored in marine environments because marine environments tend to be open water habitats with little cover available, in which diverse and large predators are present, so post-capture escape and survival is especially relevant. In contrast, the increased amount of cover available and differences in the type of predators present in freshwater habitats may favor low morphs if low morphs have a hydrodynamic advantage increasing pre-capture escape performance (Reimchen, 1992, 1994). Further, it has been hypothesized that presence of a relatively shallower body profile in many lacustrine sticklebacks, a body shape that tends to reduce hydrodynamic drag (Webb, 1984), is related to reduced overall predation pressure in these environments (Walker & Bell, 2000). Others have also shown that less derived freshwater forms tend to have deeper bodies than more derived freshwater specimens (Spoljaric & Reimchen, 2007), suggesting that adaptation to freshwater habitats is accompanied by a reduction in body depth. Alongside, the protective advantage of lateral plates is reduced in freshwater environments where avian predation dominates because birds compress fish rather than puncture them (Reimchen, 1988). So, on balance, capture avoidance becomes more important for survival and fitness in the freshwater context, in contrast to the marine environment where post-capture processes matter more (Reimchen, 1994).

Whereas most studies investigating the role of predation in producing directional selection in stickleback have focused on adult predators and adult stickleback, recent work on Salmonids, a major stickleback predator, has shown that juvenile Salmonid predators only consume juvenile stickleback, and thus do not produce a directional selective effect on *Eda* genotypes (Wasserman et al., 2021). In a large (>8000 individuals), multigenerational dataset documenting sub-lethal injuries to stickleback in a freshwater lake with both avian and piscine predators, diversifying selection was found to be an important driver of plate phenotype, and strength and direction of selection were shown to vary significantly across years (Reimchen & Bergstrom, 2023). Furthermore, this study reported that adult fish of different plate phenotypes do not exhibit the differences in injury documented for juvenile fish, and taken together these findings highlight the importance of considering life stage in such studies.

Interestingly, differences in developmental timing of plate ossification have been documented between full- and low-plate morphs (Currey et al., 2017), where ossification first begins at standard length of ∼12 mm in full-plated juveniles and much later at 16-18 mm in low-plated forms. Low-plated forms in turn attain their reduced complement of plates at approximately 22 mm, while full-plated forms do not attain a full set of plates until they reach approximately 30 mm, although at comparable sizes they have more plates developing than do low-plated morphs (Bell, 1981). Thus, at juvenile stages there is considerable variation in armour complement, and full-plated morphs are not in fact fully armoured until they reach sexual maturity.

Escape response is widely accepted as an ecologically relevant measure of maximum organismal performance, and, in fishes, escape response is defined as a fast-start, or accelerated motion, occurring in response to threatening stimuli and which may take several different forms (Domenici & Hale, 2019). Stickleback exhibit a double-bend C-start burst swimming episode during their escape response following a threatening stimulus (Taylor & McPhail, 1986). The body musculature contracts and moves the head and tail toward each other to form a C-shape which is followed by a similar contraction in the opposite direction, in a response driven by Mauthner cells (Domenici & Hale, 2019; Eaton et al., 1977). A handful of kinematic studies have investigated burst swimming responses in adult stickleback, examining the influence of various factors such as parasite load (Blake et al., 2006; Stewart et al., 2018), ecotype (Law & Blake, 1996; Morozov et al., 2018), habitat (Taylor & McPhail, 1986), and morphology (Hendry et al., 2011). These studies indicate that low-plated freshwater stickleback exhibit greater velocity and, in some cases, greater distance traveled during escape responses than do individuals from anadromous populations, both of which are known to affect survival of predator attacks (Roche et al., 2023; Walker et al., 2005). Environmental effects, however, are known to play a role in how morphology influences escape response performance in stickleback (Hendry et al., 2011), and these studies did not standardize habitat among treatment groups. In a common garden experiment where fish were raised and tested in freshwater, lateral plate number was again negatively correlated with velocity and distance traveled (Bergstrom, 2002), further supporting the hypothesis that reduced plate complement is associated with improved escape performance.

Here we investigate the relationship between body shape, performance and ontogeny through an ecomechanical lens with a focus on *Eda* genotype, in a population of stickleback recently isolated to the freshwater Lost Lagoon, British Columbia (salinity 0.08 parts per trillion (ppt)), that offers a case study of vertebrate evolution on a contemporary scale (Kozak et al., 2024). We attempt to explicitly link escape response to genotype and phenotype at juvenile and adult stages in the same population, to determine if a particular genotype conveys a performance advantage, and further to see if that advantage persists across ontogeny in both brackish and saltwater environments. This population provides a unique experimental context, because all three *Eda* genotypes and plate morphs are readily present in a single brackish habitat where gene flow is minimal, reducing the potential effects of confounding factors such as inter-habitat and inter-population variation that may influence performance comparisons. We test the hypothesis that fish with low-plated *Eda* alleles exhibit increased escape performance to avoid capture and therefore reduce predation in the absence of body armour, as previously proposed (Reimchen, 1994). Fishes experience extremely high mortality rates at early life stages, sometimes greater than 99% (Houde, 1997), and, as discussed above, there is variation among stickleback morphs in the timing of development of armour plates over ontogeny, making early life stages a powerful potential target for selective pressures. We therefore measured full body phenotype, including plate morph and body shape and size, and escape performance in a sample of adult fish from Lost Lagoon to assess differences among phenotypes and genotypes that exist in this population in the wild. We then created an F1 cross from these fish, raised them in a common brackish water environment (salinity 3-5 ppt), and investigated escape performance and phenotype at a standard length of ∼25 mm, before completion of development of lateral plates but standard lengths long enough to study escape response. This allowed us to investigate the possibility that selection is acting on juveniles rather than on adults, which are more commonly studied. Finally, we transitioned half of these fish to saltwater, and measured escape performance and phenotype again at sexual maturity. We characterized body shape phenotype in all three experimental groups using 2D geometric morphometrics rather than simple plate counts to better capture pleiotropic effects on body shape driven by *Eda* that may be evident before the completion of plate development. We predicted that possession of one or more low-plated alleles would result in better escape performance in brackish environments across ontogeny, and that there would be body shape phenotypes associated with improved performance evident at different ontogenetic stages.

## Methods

All sampling, handling, and euthanasia of stickleback was conducted in accordance with the ethical standards outlined by the Canadian Council on Animal Care and with approval of the University of Calgary Animal Care Committee (Protocol #: AC19-0016).

### Study Site and Specimen Collection

Lost Lagoon is located in Vancouver, British Columbia, Canada (49°17’44.5” N, 123°8’25.3”W), and has been isolated from the marine environment for more than 100 years (see Kozak et al., 2024). Threespine stickleback were collected from Lost Lagoon in May 2019 using minnow traps. 79 fish were collected and transported to the University of Calgary Life and Environmental Sciences Animal Resource Center. Fish were transferred into 15 US gallon Canadian Food Inspection Agency (CFIA) certified acrylic tanks and held at a density of 6-10 fish per tank, in brackish water conditions (salinity 3-5ppt), until their experimental endpoint. Each tank contained two sponge filters for aeration and water flow and two artificial aquatic plants for tank enrichment. Fish were fed blood worms (*Chironomid* sp.) until satiation once per day. Room temperature was kept at 16.0±1.0 °C and under a 16-hour light, 8-hour dark light cycle.

### Crossing and Rearing

Males and females in reproductive condition were manually crossed to produce an F1 generation consisting of three replicate families. All individuals crossed displayed the partial morph phenotype for lateral plates. Fin clips from parental fish were used to confirm heterozygosity at *Eda* for all parents following crossing (see below). The fertilized egg mass for each cross was contained in a plastic cup with a mesh bottom and placed in a separate 10 US gallon tank containing brackish water and five drops of methylene blue to prevent fungal infection.

The eggs hatched 10-13 days after crossing and F1 juveniles were then fed brine shrimp (*Artemia* sp.) until satiation twice daily until the family reached an average length of approximately 1 cm, at which time they were transitioned to chopped blood worms (*Chironomid* sp.) once daily. When juveniles reached an average standard length of ∼25 mm, they were subjected to a first round of performance trials (see below).

Following the first round of performance trials, fish were anaesthetized using 1-2 drops of clove oil (Eugenol; Sigma-Aldrich, Oakville, ON, Canada) per 0.5 litre of water under direct supervision, and visible-implant-elastomer (VIE, Northwest Marine Technology) tagged to allow individual identification throughout the experiment. VIE tags were placed on either side of the insertion point of the first dorsal spine using a 0.3 cc insulin syringe with a 29-gauge needle. The needle was administered horizontally from the anterior side of the fish 5-10mm deep such that the entirety of the needle was visible under the first layer of cells throughout the injection.

Elastomer was slowly injected as the needle was withdrawn. Each of the three F1 families was then split into four 30 US gallon tanks with 8-15 individuals per tank. All individuals were given one month to recover from tagging in brackish water conditions because VIE tagging is known to elicit a physiological response similar to that caused by the presence of a predator (Furtbauer et al., 2015), as well as an immune response in stickleback that resolves after approximately two weeks (Henrich et al., 2014). After one month of recovery time, half of each family (two tanks) was held in brackish water and half of each family (two tanks) was transitioned to saltwater (∼35 ppt over four weeks). Fish then spent another three months in these treatments before reaching sexual maturity.

### Performance Trials

Escape response performance trials conducted on wild fish and on F1 fish at the juvenile and adult timepoints used the same experimental set up and procedure. A 10 US Gallon tank was placed on a table with a cut-out so that the tank was visible from below. Tank dividers were placed in the tank to create an oval arena to prevent fish from moving out of sight of the camera and to block the experimenter and lab environment from the fish’s view. A scale bar was drawn onto the bottom of the tank to allow scaling during analysis. The tank was illuminated from both sides by 500W halogen work lights as well as from below with a 250W halogen work light. A HiSpec 1 camera (Fastec Imaging, San Diego, CA, USA) was placed below the tank and footage was recorded at 400 frames per second (fps).

All escape response trials were conducted in brackish water (3-5 ppt) except the F1 adult saltwater treatment, where trials were conducted in saltwater (35ppt) to mimic tank conditions. The experimental tank was filled with 7cm of water to limit vertical displacement during escape responses (Bergstrom, 2002; Hendry et al., 2011; Roche et al., 2023). Fish were fasted for 24 hours prior to trials and individually moved into the experimental tank where they were given 10 minutes to acclimate to the new tank with the filming lights turned off, which proved sufficient for recovery from handling stress in pilot trials. After 10 minutes, the camera and lights were turned on and an escape response was elicited from the fish by manually tapping a probe directly behind the fish to simulate a predator (Bergstrom, 2002). This was repeated a minimum of five times for each fish with the goal of obtaining three complete, uninhibited, and fully in-frame escape responses. Five trials were used to ensure performance maximums were captured (Nelson & Claireaux, 2011). Fish were given five minutes of rest time between replicates to allow for recovery. Lights were turned off between trials to prevent water temperature from rising. Water was refreshed between individuals to maintain a consistent temperature. In total, three successful escape response trials were collected for 49 wild adult fish, 85 individuals from two families at the juvenile timepoint, and 119 individuals from three families at the adult timepoint (85 individuals from the juvenile timepoint and an additional 34 individuals from a third family) to provide additional statistical power considering the additional complexity of the salinity treatments for a total of 612 trials for analysis.

Escape response videos were labeled with three distinct points in every frame using a deep neural network created with the Python package DeepLabCut v.2.1.6.4 (Mathis et al., 2018; Nath et al., 2019) that allows high throughput markerless pose estimation for very large datasets (see Supplementary Materials for details of this procedure). The anterior tip of the snout, the midpoint between the pectoral fins, and the caudal peduncle were labeled using this method (Bergstrom, 2002). Labeled videos were reviewed and the first three videos for each individual that met quality criteria were selected for analysis. Videos did not meet criteria at this step if: the individual did not respond to stimuli or responded with a gliding movement not classified as a C-start; the individual swam out of frame or the entirety of the response was not captured; the response was inhibited by running into the arena wall; or the network labels tracked poorly.

Tracks of label coordinates were smoothed with a three-point moving average. The smoothed data were then used to calculate the average maximum velocity achieved during the escape response; the average curvature coefficient, defined as the smallest distance between the nose and the tail during the response divided by standard length (Bergstrom, 2002), and the average total distance traveled during Stages 1 and 2 of the double bend escape response. The beginning of the response was identified as the last frame after the escape stimulus and before the angle between the three points on the fish changed by at least 0.5 degrees. The end of the escape response was identified as the second time the points realigned to capture a double bend C-start (Domenici & Hale, 2019). In some cases, the three points did not realign a second time because the fish either contracted in the same direction a second time before returning to center or the fish stopped moving before finishing the second contraction. In these cases, the maximum angle (approaching 180 degrees) achieved after the second contraction was used as the end point for the response. Measurements of standard length (SL) were taken from the anterior tip of the snout to the caudal peduncle, using calipers on anaesthetized or euthanized fish.

### Euthanasia and Preservation

Once all trials were complete, individuals were euthanized using 2-4 drops of clove oil (Eugenol; Sigma-Aldrich, Oakville, ON, Canada) per 0.5 L of water. Caudal and pectoral fins were first removed and placed in 95% ethanol. Specimens were then fixed in 10% neutral buffered formalin for a minimum of two weeks before being moved to 70% ethanol for long-term storage.

### Sex and *Eda* genotyping

DNA was exacted from fin clips following a modified protocol with a Qiagen DNeasy Blood and Tissue kit (Barry, 2019). Sex genotyping was performed using Isocitrate dehydrogenase (*Idh*) which is deterministic of sex in threespine stickleback (Peichel et al., 2004). *Eda* was genotyped using a diagnostic intel marker in intron one (stn382) (Colosimo et al., 2005; Le Rouzic et al., 2011). Sex and *Eda* genotyping were performed at the Molecular Biology Facility at the University of Alberta.

### 2D Geometric Morphometric Analysis

Following each set of escape response trials, a DinoLite Edge digital microscope camera (Dunwell Tech Inc., Torrance, CA, USA) was used to photograph each fish. Photographs were taken from left lateral views with a scale bar. Juvenile F1 fish were photographed alive immediately after mild anaesthesia and VIE tagging. Wild fish and adult F1 fish were euthanized prior to photography, following their final escape response trial.

Image files of the left side of the fish were compiled into TPS files using tpsUtil32 v.1.79 (Rohlf, 2019) and landmarked in tpsDig2w32 v.2.31 (Rohlf, 2018) using a set of 25 landmarks (Fig. 1) adapted from those of Bjærke et al. (Bjærke et al., 2010) with the goal of capturing body shape variation across ontogeny. The same landmark set was used for wild fish and F1 fish at both juvenile and adult timepoints. To evaluate accuracy in landmark placement, particularly given the difficulty of landmarking juvenile fish, a landmark trial was conducted using 20 individuals from each of the wild, juvenile F1, and adult F1 datasets. The landmark set was applied to the lateral left view of each fish three separate times, and Procrustes analysis of variance was used to confirm that landmark placement was not a significant contributor to the variation in the dataset. We then proceeded with the main analysis using a single landmark set for each individual fish at each timepoint.

**Figure 1.**
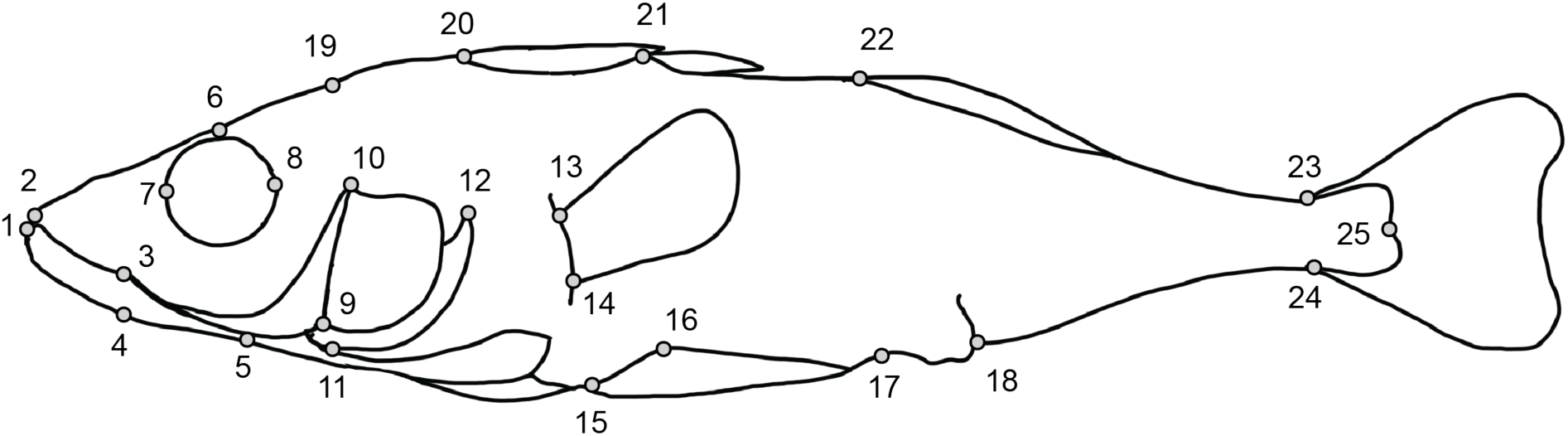
2DLandmarks collected for this study, adapted from Bjærke, et al. (Bjærke, et al., 2010). 1: anterior extent of dentary; 2: anterior extent of maxilla; 3: posterior extent of mouth; 4: posterior extent of dentary; 5: inferior-most point below orbit; 6: superior-most point of head above orbit; 7: anterior extent of orbit; 8: posterior extent of orbit; 9: ventral-most point of operculum; 10: anterodorsal tip of operculum; 11:anterior extent of suboperculum; 12: superior extent of suboperculum; 13: superior insertion of pectoral fin; 14: inferior insertion of pectoral fin; 15: posterior extent of ectocoracoid; 16: superior extent of pelvic plate; 17: posterior extent of pelvic plate; 18: anterior insertion of anal fin; 19: anterior extent of supraoccipital; 20: anterior insertion of first dorsal spine; 21: anterior insertion of second dorsal spine; 22: anterior insertion of dorsal fin; 23: dorsal insertion of caudal fin; 24: ventral insertion of caudal fin; 25: posterior extent of caudal peduncle.

Landmark data for wild fish, F1 fish at the juvenile timepoint, and F1 fish at the adult timepoint were analyzed separately using geomorph v.4.0.10 (Adams et al., 2025; Baken et al., 2021) and RRPP v.2.1.2 (Collyer & Adams, 2024) in R 4.3.3 (R Core Team, 2025) running in RStudio v.2024.12.1 (Posit Software, 2025). For each group, landmark coordinates were combined into a single TPS file and read in as a matrix. Data were Procrustes-transformed to remove the effects of translation, rotation, and scale (Dryden & Mardia, 1998) and then converted into a geomorph data frame, and additional variables of interest were added to the geomorph data frame. In the wild dataset this included sex and *Eda* genotypes. In the F1 juvenile dataset this included sex, family, and *Eda* genotype and in the F1 adult dataset this included sex, family, *Eda* genotype, and salinity treatment. Principal components analysis (PCA) was performed separately on each set of Procrustes-transformed landmark coordinates to reduce dimensionality and visualize major axes of shape variation. High-dimensional Procrustes RRPP-ANOVA (Collyer et al., 2015) was applied to the transformed coordinates, including the natural log of the centroid size of the landmark configuration (logCsize) which functions here as a proxy for size (Klingenberg, 2016) as a covariate, to assess differences in shape. Backward selection was used to reduce these models where appropriate. Finally, the first principal component of the shape analysis was regressed on logCsize within each group, to obtain a size-corrected index of shape for that group, and this variable was considered as a body shape covariate in the escape response performance analyses described below.

### Kinematic Analysis

Analysis of variance was used to determine the effects of sex, *Eda* genotype, family and salinity treatment (for F1 adults only) on standard length. Linear models were then constructed for each performance variable: maximum velocity, curvature coefficient, and total distance traveled. Total distance traveled was natural-log transformed to produce a normal distribution. Analysis of covariance, with standard length as a covariate to control for the effects of standard length on kinematic variables (except in the case of curvature coefficient, which is calculated as maximum of curvature divided by standard length), was used to assess the contribution of sex, *Eda* genotype, family and salinity treatment (for F1 adults only) for each set of trials. This analysis was repeated with the body shape covariate obtained as described above in place of standard length to determine if size or shape was a better predictor of performance for each variable. Backward selection was once again used to reduce these models where appropriate. Escape performance analyses were conducted in R 4.3.3 (R Core Team, 2025) running in RStudio v.2024.12.1 (Posit Software, 2025), and all plots were produced using ggplot2 v.3.5.1 (Wickham, 2016).

## Results

### *Eda* Genotypes and Standard Lengths

Figure 2 shows *Eda* genotypes by sex for the wild caught (Fig 2A) and F1 adult (Fig 2B) fish. In the wild caught group (n= 49), there were 18 individuals homozygous for the full-plated allele, 8 homozygous for the low-plated allele, and 23 heterozygotes. Among the F1 adult fish there was an approximately Mendelian distribution of genotypes with 32 individuals homozygous for the full-plated allele, 29 homozygous for the low-plated allele, and 58 heterozygotes (n = 119), and in the subset of these used for the juvenile analysis (n = 85), there were 21 individuals homozygous for the full-plated allele, 24 homozygous for the low-plated allele, and 40 heterozygotes. Standard lengths for each experimental group are plotted in Figure 3. For the wild group (Fig 3A), sex (p = 0.0396) was the only significant contributor to variation in size. Similarly, among the F1 juveniles (Fig 3B), sex (p = 0.0017) was the only significant contributor to variation in size. Among the F1 adults, (Fig 3C), *Eda* genotype (p = 0.0016) and salinity treatment (p < 0.0001) both contributed to variation in size, and there was a significant interaction between salinity and sex (p = 0.0425) which is elaborated on further below.

**Figure 2.**
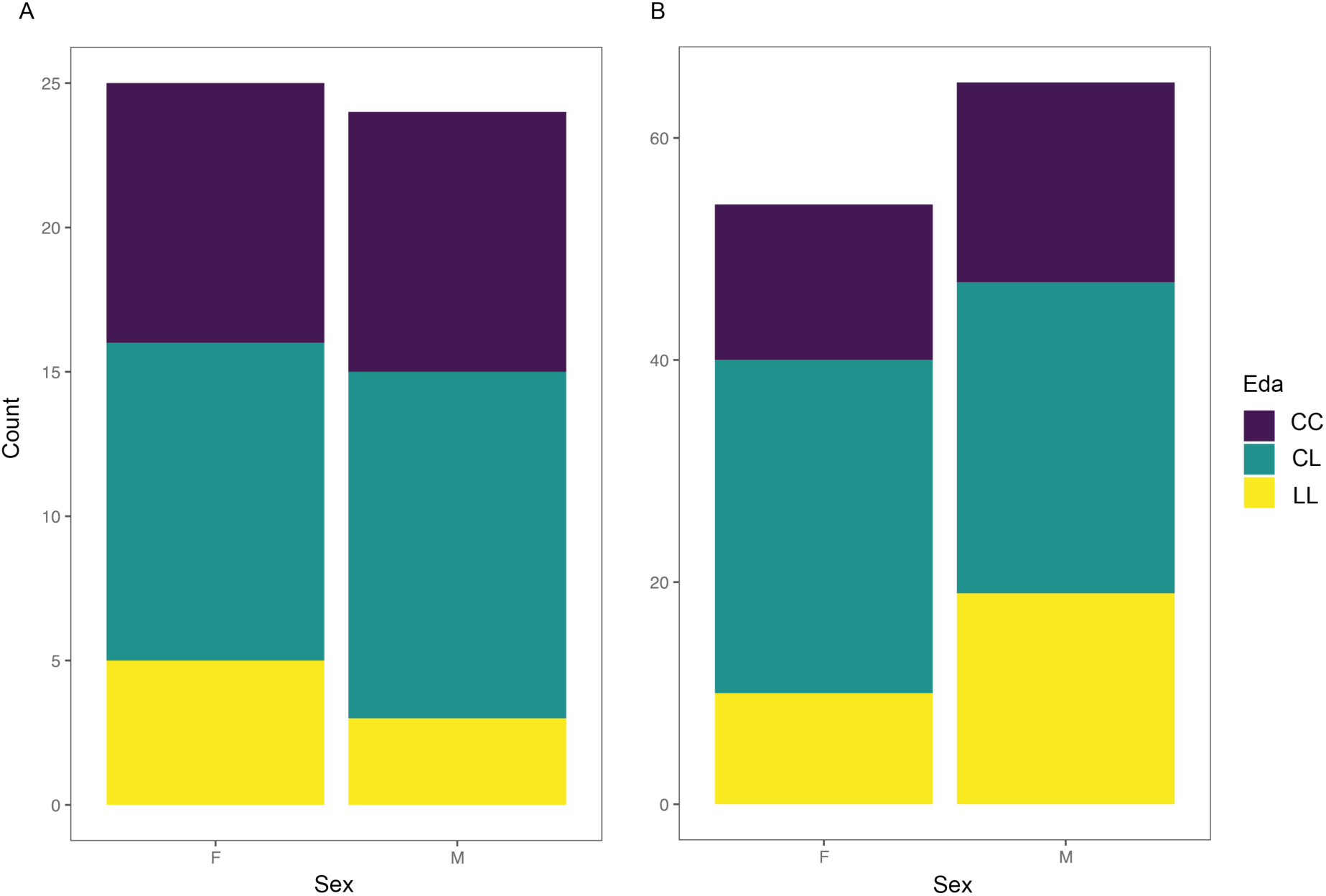
*Eda* genotypes by sex for wild caught (A; n = 49) threespine sticklebacks in Lost Lagoon, BC and F1 adult (B; n = 119) threespine sticklebacks raised in the laboratory. A subset of the F1 adult group was also studied at a juvenile timepoint (n = 85). Abbreviations: C: full-plated allele; Eda: Ectodysplasin; F: female; L: low-plated allele; M: male.

**Figure 3.**
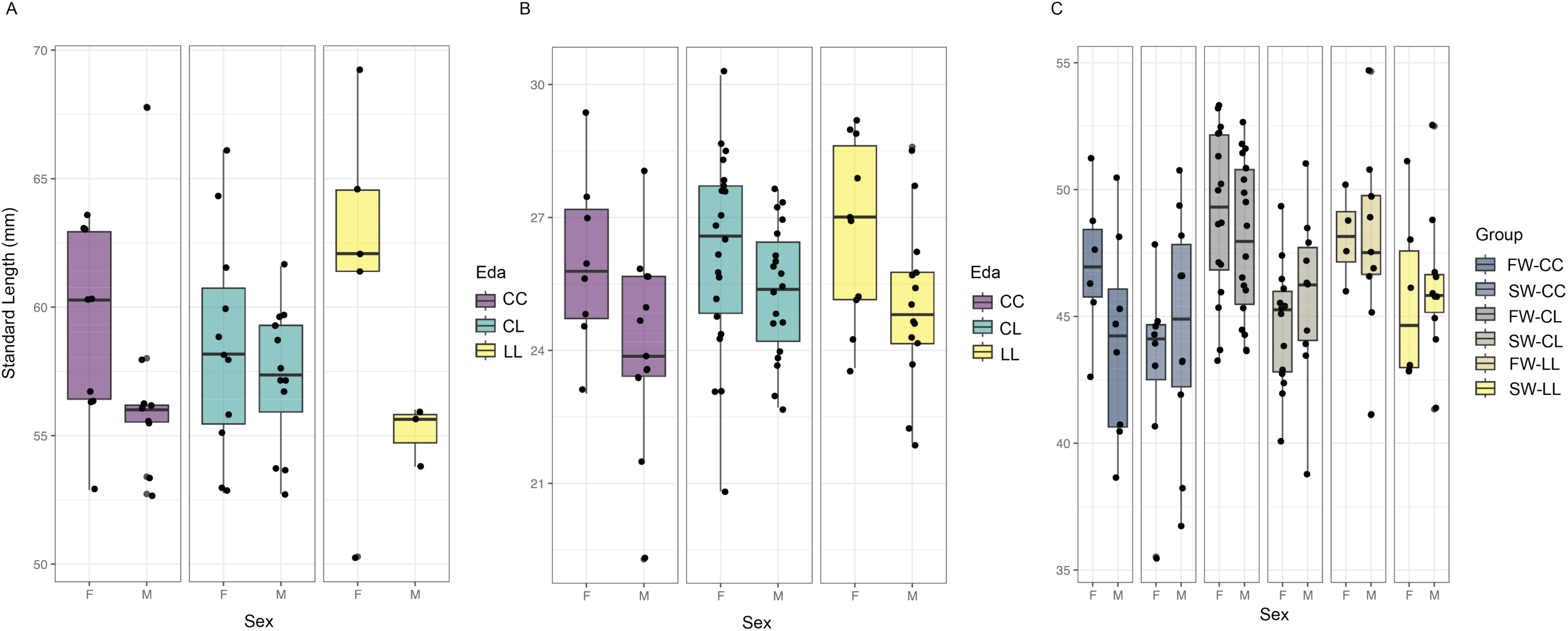
Distributions of standard lengths for wild caught (A), F1 juvenile (B), and F1 adult (C) threespine sticklebacks used in escape performance trials. Abbreviations: Abbreviations: C: full-plated allele; Eda: Ectodysplasin; F: female; FW: freshwater; L: low-plated allele; M: male; SW: saltwater.

### 2D Geometric Morphometric Analysis

Principal components analysis of the wild-caught adult fish did not reveal any substantive clustering of individuals by genotype (Fig 4A). Females tended to cluster on the positive end of PC1 while males clustered on the negative end, regardless of genotype, suggesting a strong contribution of overall size to this component, which explained 27.0% of the variance in the dataset. This axis captured variation in body depth and elongation, with stouter individuals at one end and more streamlined individuals at the other. PC2, accounting for 17.9% of the variance did not differentiate among individuals by genotype or sex. This axis captured minor variation in skull depth, but in general the shape changes along this axis were ill-defined. LogCsize was a significant contributor to shape (p = 0.0003), and after accounting for size, sex (p = 0.0010) was a significant contributor to variation among wild-caught adults, whereas *Eda* genotype was not, indicating sexual dimorphism in shape that is unrelated to *Eda* genotype.

**Figure 4A.**
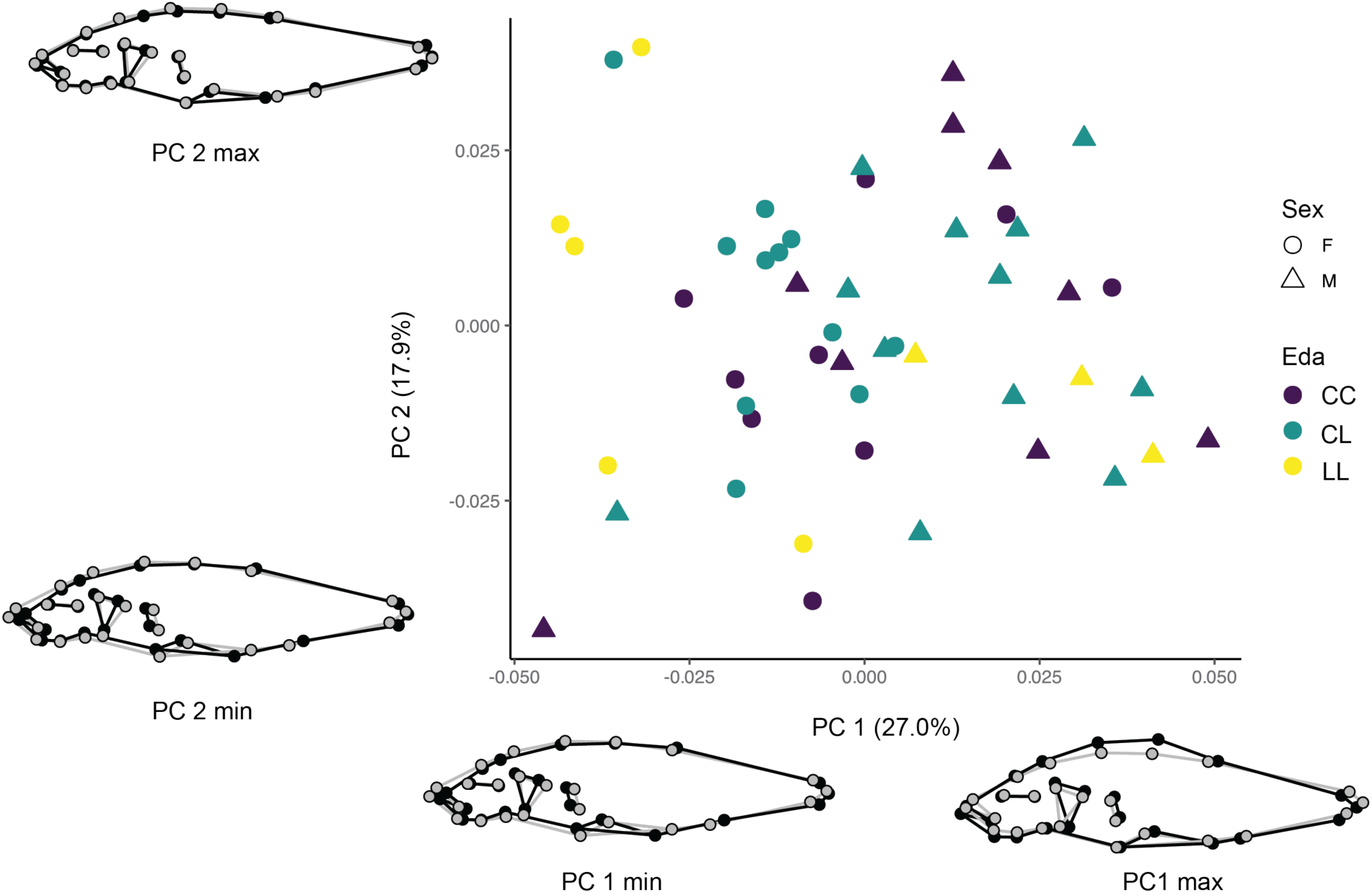
Principal components analysis of body shape in wild caught threespine sticklebacks in Lost Lagoon, BC. Grey wireframe indicates mean shape for each landmark configuration, black wireframe indicates configuration at each end of the indicated axis. Abbreviations: C: full-plated allele; Eda: *Ectodysplasin*; F: female; L: low-plated allele; M: male.

Principal components analysis of the F1 juvenile fish again revealed no discernable clustering relative to *Eda* genotype, and in contrast to the wild-caught sample, there was no pattern associated with sex (Fig 4B). PC1 explained 27.3% of the variation in the dataset, and again represented variation in body depth and elongation, with stouter individuals at one end and more streamlined individuals at the other. PC2 explained 12.6% of the variation, capturing variation in body depth. Once again logCsize was a significant contributor to shape (p = 0.0364), and there were no other significant contributing factors.

**Figure 4B.**
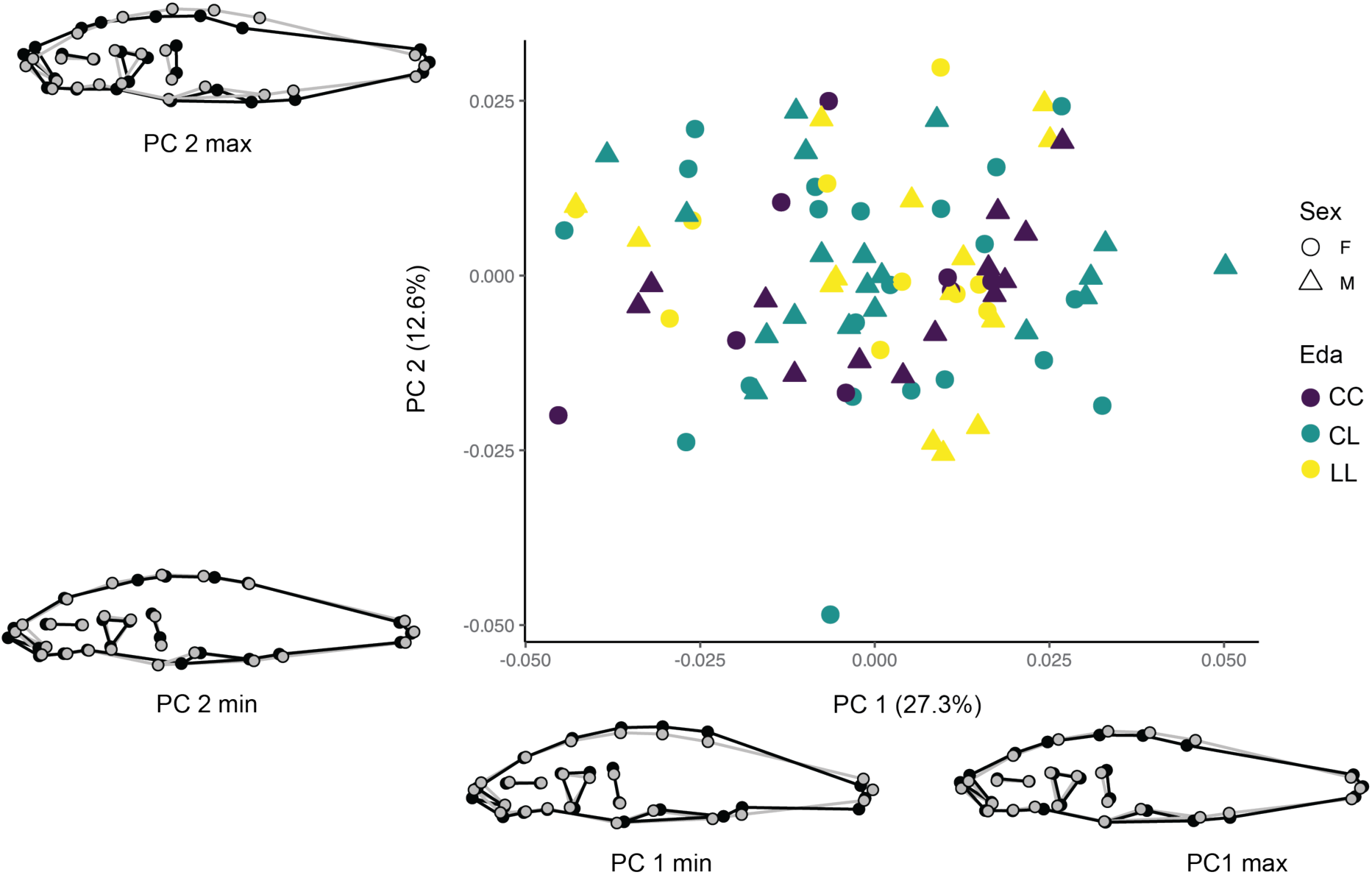
Principal components analysis of body shape in F1 juvenile threespine sticklebacks in Lost Lagoon, BC. Grey wireframe indicates mean shape for each landmark configuration, black wireframe indicates configuration at each end of the indicated axis. Abbreviations: C: full-plated allele; Eda: *Ectodysplasin*; F: female; L: low-plated allele; M: male.

Principal components analysis of the F1 adults did not produce any definitive clustering by *Eda* genotype or salinity treatment but did indicate, similar to the scenario in the wild-caught adults, a pronounced sexually dimorphic aspect to shape on PC1, which explained 24.0% of the variance in the dataset, and toward the positive end of which females of all three genotypes and both treatments tended to cluster (Fig 4C). Like the results for the wild-caught adults, this axis captured variation in body depth and elongation, differentiating stouter individuals from more streamlined specimens. This axis also captured variation in operculum area and relative placement of the pectoral fin. PC2 explained 13.6% of the variance in the dataset and did not differentiate by treatment, sex, or *Eda* genotype. This axis represented variation in body depth. Once again logCSize was a significant contributor to shape (p = 0.0001), and sex (p = 0.0001), treatment (p = 0.0012) and *Eda* genotype (0.0453) all contributed significantly to shape variation in this sample after accounting for size.

**Figure 4C.**
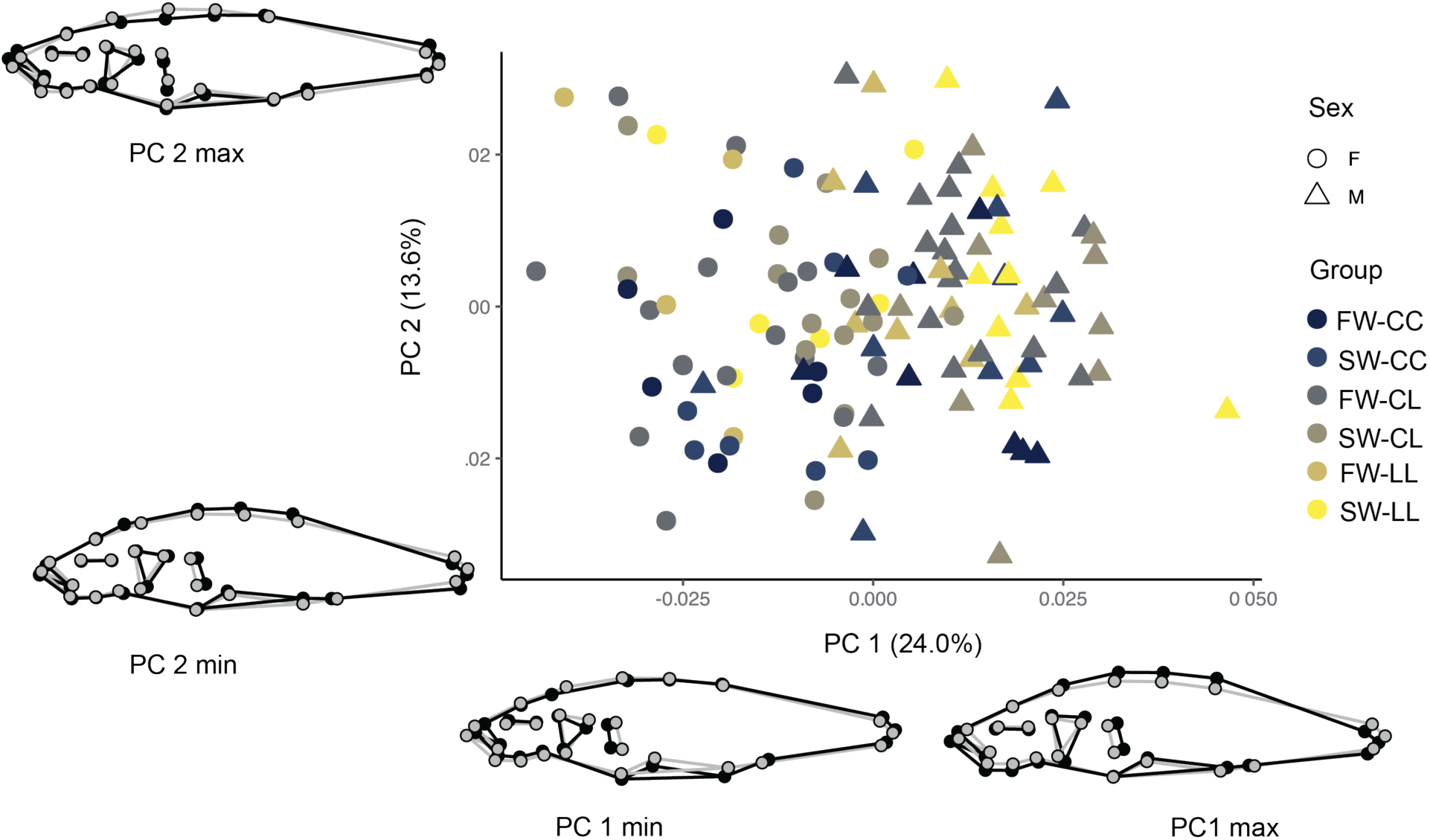
Principal components analysis of body shape in F1 adult threespine sticklebacks in Lost Lagoon, BC. Grey wireframe indicates mean shape for each landmark configuration, black wireframe indicates configuration at each end of the indicated axis. Abbreviations: C: full-plated allele; F: female; FW: freshwater; L: low-plated allele; M: male; PC: principal component; SW: saltwater.

### Kinematic Analysis

Escape response variables for the wild-caught adult fish are plotted in Figure 5. Statistical analysis of these results revealed no significant differences between sexes for maximum velocity or curvature coefficient, and length did not contribute to variation in either of these performance measures. Standard length was a significant contributor to total distance traveled (p = 0.0177), with longer fish traveling further. *Eda* genotype did not significantly affect any aspect of escape performance, and there were no significant interactions among variables in any of these models. Wild-caught females were significantly longer than males (p = 0.0353). *Eda* genotype had no significant impact on standard length. Body shape did not significantly influence any aspect of escape performance.

**Figure 5.**
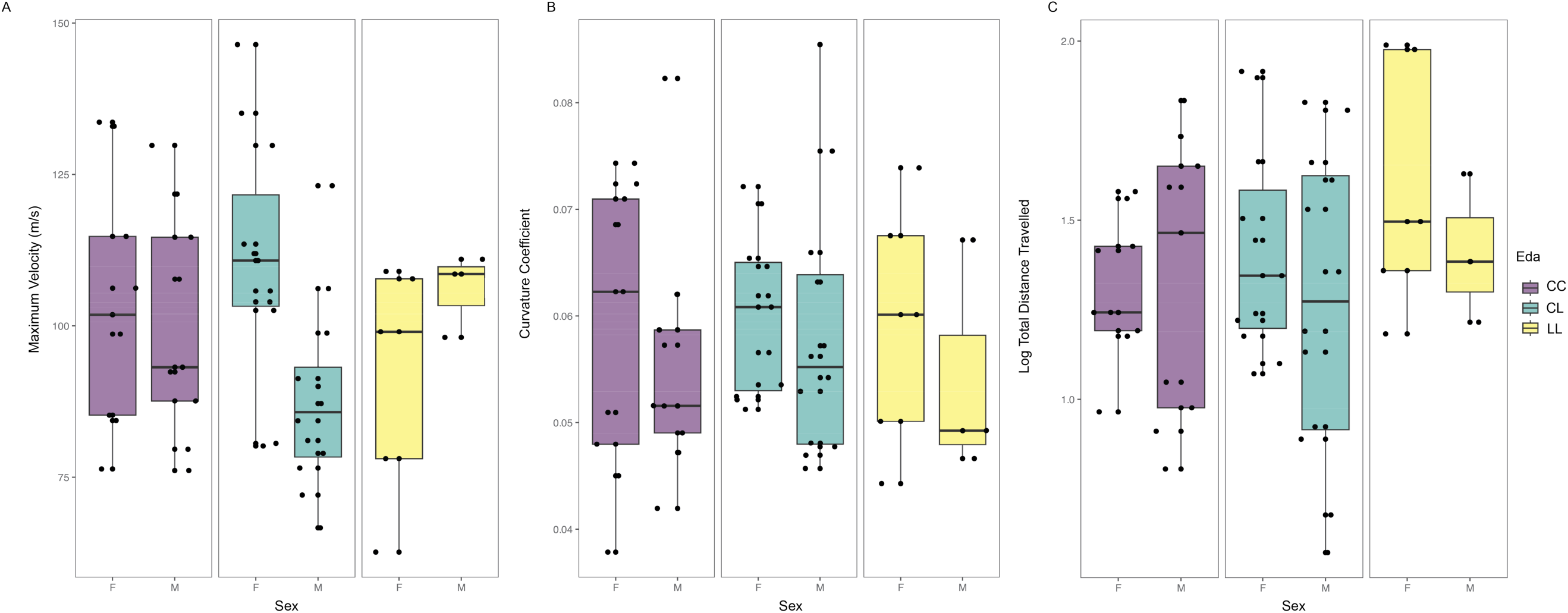
Escape response variables measured in this study for wild caught adult threespine sticklebacks. A) Maximum velocity; B) Curvature coefficient; C) Natural log of total distance traveled. Abbreviations: C: full-plated allele; Eda: *Ectodysplasin*; F: female; L: low-plated allele; M: male.

Escape response variables for the wild-caught adult fish are plotted in Figure 6. Among F1 juvenile fish, standard length was significantly associated with maximum velocity (p = 0.0019) and total distance traveled (p = 0.0004), with longer fish reaching a greater maximum velocity and greater total distance traveled, and there was also a significant effect of family on maximum velocity (p = 0.0015) and curvature coefficient (p= 0.0004). *Eda* genotype did not significantly affect any aspect of escape performance, and there were no significant interactions among variables in any of these models. Once again females were significantly longer than males (p = 0.0020), but *Eda* genotype had no significant impact on standard length. Body shape significantly influenced maximum velocity (p = 0.0034) and curvature coefficient (p = 0.0007), but not total distance traveled.

**Figure 6.**
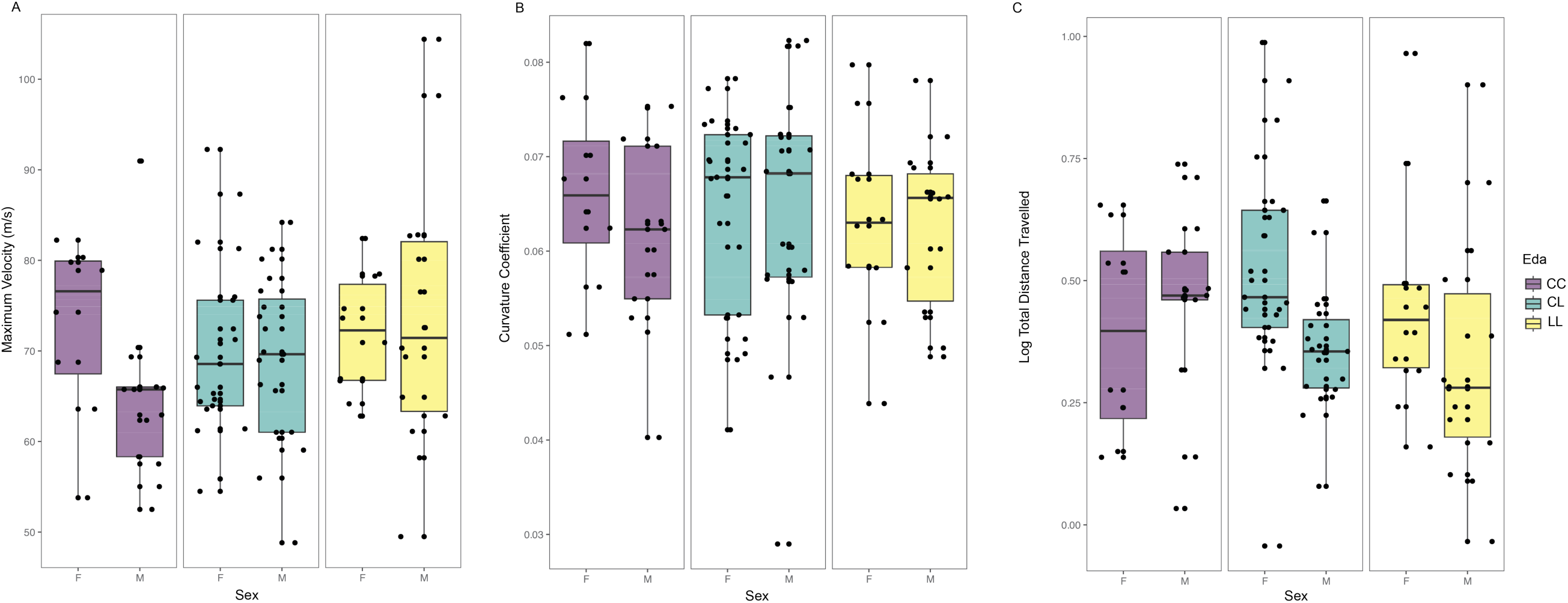
Escape response variables measured in this study for F1 juvenile threespine sticklebacks. A) Maximum velocity; B) Curvature coefficient; C) Natural log of total distance traveled. Abbreviations: C: full-plated allele; Eda: *Ectodysplasin*; F: female; L: low-plated allele; M: male.

Escape response variables for the wild-caught adult fish are plotted in Figure 7. Among F1 adult fish, standard length was significantly associated with maximum velocity (p = 0.0093), whereas no other variables produced significant effects. Standard length (p = 0.0367), sex (p = 0.0003), and family (p = 0.0013) all had significant effects on curvature coefficient. Neither *Eda* genotype nor salinity treatment significantly affected maximum velocity or curvature coefficient, and there were no significant interactions among variables in any of these models.

**Figure 7.**
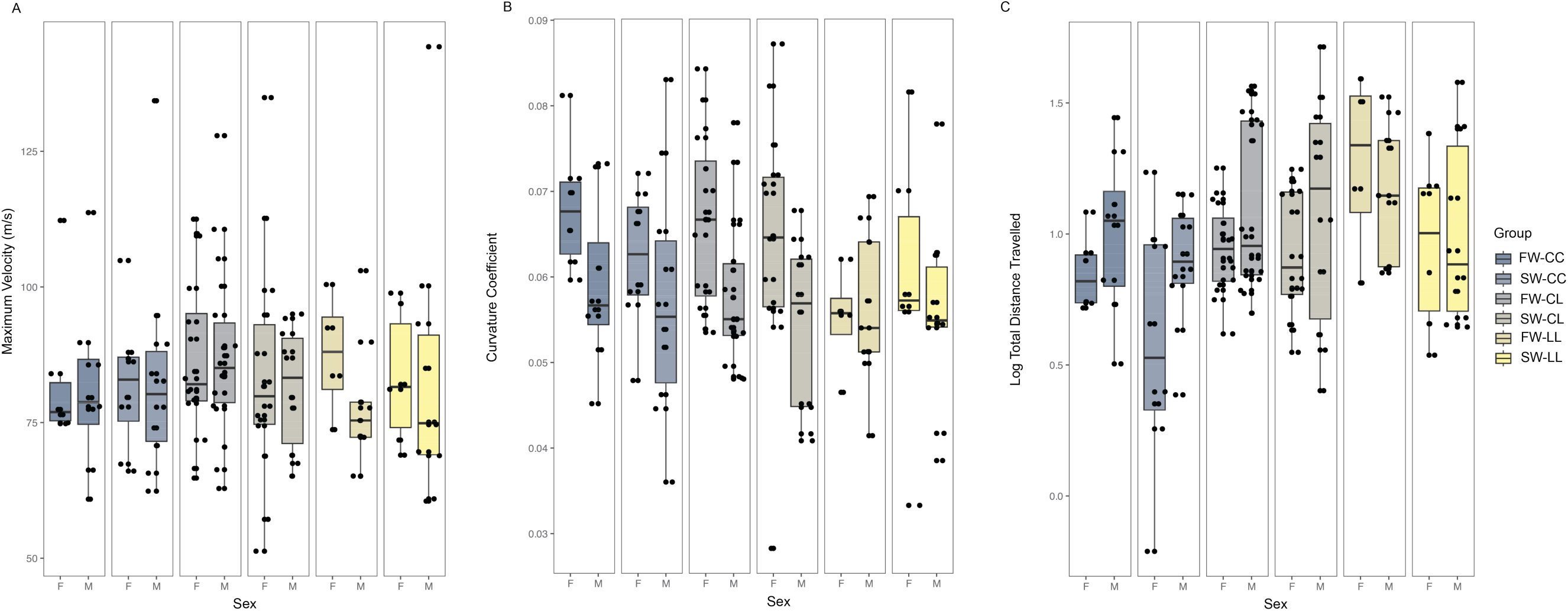
Escape response variables measured in this study for F1 adult threespine sticklebacks. A) Maximum velocity; B) Curvature coefficient; C) Natural log of total distance traveled. Abbreviations: C: full-plated allele; F: female; FW: freshwater; L: low-plated allele; M: male; PC: principal component; SW: saltwater.

Standard length (p = 0.0005), *Eda* genotype (p = 0.0331), and sex (p = 0.0100) all had significant effects on total distance traveled (Fig 7C), and there was a significant three-way interaction among standard length, *Eda* genotype, and sex (p = 0.0046). Salinity treatment had no significant effect on total distance traveled. In this group, *Eda* genotype (p = 0.0019) and salinity treatment (p < 0.0001) had significant effects on standard length, where fish with one low-plated allele (p = 0.0019) or two low-plated alleles (p = 0.0167) traveled further than those homozygous for the full-plated allele, and fish raised in fresh water (p < 0.0001) were longer than those raised in salt water. In this sample, sex was not a significant contributor to variation in standard length. Body shape significantly influenced coefficient of curvature (p = 0.0202), but did not influence maximum velocity or total distance traveled in this group.

## Discussion

In this study, we sought to link escape response performance to genotype and phenotype across ontogenetic stages within a single population, to determine if *Eda* genotype conveys a performance advantage in either brackish or saltwater environments. We predicted that possession of one or more low-plated alleles would result in better escape performance in brackish environments across ontogeny, and that *Eda* genotype would result in variation in shape phenotype that would be predictive of escape performance before the completion of lateral plate development. We found that *Eda* genotype does not directly impact maximum velocity or curvature coefficient but is linked to an increase in total distance traveled in F1 adults. We further showed that *Eda* genotype impacts growth, producing variation in size which in turn impacts escape performance variables across age groups, with a particular focus on total distance traveled. Although we showed a statistical association between *Eda* genotype and shape, this variation was subtle and is confounded by sexual dimorphism as shown by others, and further study is required to better tease out the relationship between *Eda-*associated shape variation and performance. The presence of at least one low-plated allele appears to have conferred a growth advantage in this experiment, as did a freshwater rearing environment, suggesting that pleiotropic effects of *Eda* on skeletal development may be modulating anti-predator responses before plate phenotype is fully expressed.

Altered predation regimes are hypothesized to impose selection pressures that favour a low-plated phenotype on threespine stickleback in freshwater (Barrett, 2010; Leinonen et al., 2011; Reimchen, 1994). Many vertebrate and invertebrate taxa prey on stickleback, ranging from predatory fishes and birds to freshwater macroinvertebrates (reviewed in (Reimchen, 1994)). In particular, the ability to escape from predators before capture should be selectively advantageous for stickleback in freshwater environments, where the low-plated *Eda* allele becomes dominant and results in reduced body armour and reduced ability to survive a predation attempt once captured (Reimchen, 1992, 1994). This contrasts with stickleback in saltwater environments, where the number and variety of predators is much larger, and where the prevalence of full-plated *Eda* alleles produces individuals with a full complement of body armour that should be better able to physically withstand a predation event upon capture.

Following previous work on stickleback escape response and burst swimming (Bergstrom, 2002; Law & Blake, 1996; Morozov et al., 2018; Taylor & McPhail, 1986), we selected three kinematic variables to describe escape performance: curvature coefficient, maximum velocity, and total distance traveled. These variables were selected to provide an overview of the components of this response that are likely to be influenced by reduction in lateral plates associated with low-plated *Eda* alleles. Our study differs from previous work by including fish with all three *Eda* genotypes obtained from a single habitat with little to no gene flow (Kozak et al., 2024) and thus controlling for variation induced by both genetic background and habitat using a naturally occurring population, and by taking a longitudinal approach to escape performance testing to investigate variation in these variables and their relationship to genotype and salinity over time.

As expected based on previous work (e.g. (Bainbridge, 1958; Blake, 2004; Domenici, 2001) we showed that body size, measured here as standard length, contributes to escape response performance across ontogeny and is more important in this experiment than body shape, producing significant variation in many more kinematic analyses than did body shape. Here we showed that total distance traveled was significantly positively correlated with standard length across ontogeny and in the wild sample. In F1 juveniles, it was also significantly positively correlated with maximum velocity. In the wild and juvenile groups, females were significantly longer than males, but once we accounted for variation induced by standard length, we did not observe an effect of sex on any kinematic variable in either of these groups. We also showed that the presence of at least one low-plated *Eda* allele was associated with larger body size in our F1 adult group, as was rearing in fresh water (Fig 3C), and here the positive effect of standard length on total distance traveled was also observed for maximum velocity and curvature coefficient, indicating a more global impact of body size on performance in this group.

After accounting for the effects of size, our escape response results are only partially congruent with previous work. We found evidence for greater distance traveled by F1 adults with at least one low-plated allele (Fig 7C), which agrees with some previous work (Bergstrom, 2002; Taylor & McPhail, 1986). We did not observe any differences in maximum velocity among *Eda* genotypes at any stage, whereas others have reported that adult low-plated morphs are capable of greater maximum velocity (Bergstrom, 2002; Law & Blake, 1996; Morozov et al., 2018; Taylor & McPhail, 1986). Low-plated morphs have been found to be capable of greater maxima of curvature, indicating greater body flexibility, in some cases (Taylor & McPhail, 1986) but not others (Bergstrom, 2002), whereas we report no effect of *Eda* genotype on curvature coefficient but rather an effect of family on this variable in our F1 group that persists across ontogeny. We also found an effect of family on maximum velocity in juveniles, indicating the potential for considerable variation in escape response at early ontogenetic timepoints. It is possible that inherited aspects of muscle physiology and metabolism may be producing these transient family effects, or that variation in escape response is reduced in adults in comparison to juveniles, as repetition and learning produce a more homogenous response. Further work is required to better understand these effects.

Rather than relying on plate phenotype, we assessed full-body phenotype using geometric morphometrics to better understand pleiotropic effects of *Eda* on body shape beyond lateral plates and at a life stage at which lateral plates are incompletely developed. We showed that size is a significant contributor to body shape across ontogeny (Klingenberg, 1998) and we also found significant effects of sex on shape in both wild and F1 adult samples after accounting for size, as has been demonstrated by many others (e.g. (Aguirre & Akinpelu, 2010; Albert et al., 2008; Kitano et al., 2007; Schutz et al., 2022). In addition to sex, we found significant effects of *Eda* genotype and rearing conditions on shape in the F1 adults, although the nature of these shape effects was complex and confounded by the observed sexual dimorphism, which appears to reflect a stouter body shape in adult females. We were not able to replicate the results of others who have shown that freshwater forms tend to have more gracile body shapes (Spoljaric & Reimchen, 2007; Walker & Bell, 2000), but our finding that the effects of *Eda* on body shape are interrelated with the effects of sex agrees with previous work (Albert et al., 2008). Our shape findings are concordant with our escape response results, however, which indicated that sex and *Eda* genotype were significant contributors to variation in total distance traveled, and further that our body shape index showed a significant interaction with *Eda* genotype in structuring variation in total distance traveled in F1 adults. We also found evidence for effects of shape on coefficient of curvature and maximum velocity in juveniles, although these shapes were not significantly associated with *Eda* genotype at this time point. These results imply that altered body shapes may confer an advantage for some individuals in avoiding capture, even before plate complements are fully developed among juveniles within a population; however, the interplay between *Eda* genotype and sex requires further investigation to clarify the nature of this advantage.

Taken together, these results agree with those of previous studies which suggest that *Eda* genotype impacts growth, has pleiotropic effects on shape and, in this case, produces larger size associated with the low-plated phenotype in fresh water (Albert et al., 2008). As discussed above, the low-plated phenotype has been shown to be adaptive in freshwater environments (Barrett, 2010), although the precise mechanism for this advantage is not well understood. Here we show some evidence for the hypothesis that osmoregulatory genes near *Eda* may be implicated (Heuts, 1947), based on our identification of an apparent interaction between *Eda* genotype and rearing environment. Our results also support the hypothesis that there may be a trade-off between plate production and growth (Marchinko & Schluter, 2007) in freshwater that results in a growth advantage for low-plated forms. We did not continue our experiment long enough to assess longer term survival (Barrett et al., 2010) and did not include temperature variation as a variable in this experiment, as suggested by others (Kanbe et al., 2023; Smith et al., 2021). Our results are consistent with the hypothesis that *Eda* genotype may be modulating escape response prior to plate development, but our results more strongly suggest that the growth advantage conferred by low-plated *Eda* alleles in freshwater is more important for fitness. Whereas this advantage may be beneficial for escaping predation by producing heightened escape responses before plates are fully developed, escape response variables themselves may not necessarily be direct targets of selection on *Eda* genotype given that the ability to grow relatively more rapidly than conspecifics is likely to confer many other fitness advantages, such as the production of larger numbers of eggs by females (Schluter et al., 2021). Our results suggest that females with at least one low-plated allele tend to be larger in freshwater environments (Fig 3C), which further supports this hypothesis.

The effects we report here are relatively small, and it is thus important to note that larger sample sizes are likely required to more fully tease apart the subtle impacts of different variables on performance. For example, further studies with larger sample sizes may help to clarify if there is a link between specific body shapes, *Eda* genotype and performance at juvenile timepoints, where our results indicated confounding effects of family-level variation that may be more important that *Eda* genotype early in ontogeny. Our work was also limited by the constraints of the testing apparatus we used to measure escape response variables, where fish were required to reach a minimum size before being suitable for this study. The ability to collect escape response data from earlier timepoints, and further to continue the study for multiple seasons, would afford the opportunity to better characterize the ontogeny of escape response and its relationship to body shape and plate formation across the lifespan. Finally, we note that we did recover family effects in some analyses, suggesting that among-individual variation is high for the variables we measured, particularly at juvenile timepoints. This variability is interesting for future studies of selective advantage, but tackling this question was beyond the scope of this study.

In conclusion, we report a strong positive link between standard length and escape response performance in threespine stickleback adapting to freshwater across the lifespan and further provide support for previous studies indicating that the low-plated *Eda* genotype produces a growth advantage in freshwater environments that appears to be more meaningful for females. We further found effects of shape on some performance variables at juvenile stages, suggesting that body shape is important for escape response before armour plates are fully formed.

## Acknowledgements

We are grateful to Doug Syme, the members of the Lucas, Rogers, and Jamniczky labs, and the staff of the Bamfield Marine Sciences Centre, whose support was critical to the completion of this project. The University of Calgary is situated within the traditional territories of the people of the Treaty 7 region in Southern Alberta, which includes the Blackfoot Confederacy (comprising the Siksika, Piikani, and Kainai First Nations), as well as the Tsuut’ina First Nation, and the Stoney Nakoda (including the Chiniki, Bearspaw, and Goodstoney First Nations). The City of Calgary is also home to Métis Nation of Alberta, Districts 5 and 6. We acknowledge and respect the traditional caretakers of the lands in which we collected samples and performed this work, including the Musqueam, Squamish and Tsleil Waututh nations.

## Contributions

AMK developed ideas, collected and analyzed data, contributed to manuscript preparation.

MC analyzed data, contributed to manuscript preparation.

SMR developed ideas, contributed to manuscript preparation, secured funding.

KNL developed ideas, analyzed data, contributed to manuscript preparation.

HAJ developed ideas, analyzed data, contributed to manuscript preparation, secured funding.

## Data Availability

Raw data and code will be archived to Dryad upon acceptance.

