## Supplemental Methods for "Ontogeny of escape response and body shape in Threespine Stickleback (*Gasterosteus aculeatus* L.)"

**Running title**: stickleback escape response ontogeny

Aspen M. Kozak[1], Michael Chung[1], Sean M. Rogers[2], Kelsey N. Lucas[2,a], Heather A. Jamniczky[1,*,a]

1. Department of Cell Biology & Anatomy, University of Calgary, Calgary, Alberta, Canada
2. Department of Biological Sciences, University of Calgary, Calgary, Alberta, Canada

[a] Contributed equally

[*] **Corresponding author:**

**Heather Jamniczky**

Cumming School of Medicine

University of Calgary

3330 Hospital Dr NW

Calgary AB T2N 4N1

**SUPPLEMENTARY MATERIAL**

**Markerless Pose Estimation**

Escape response videos collected for this study were labeled with three distinct points in every frame using a deep neural network created with the Python package DeepLabCut v.2.1.6.4 (Mathis et al., 2018; Nath et al., 2019) that allows high throughput markerless pose estimation for very large datasets. The details of this procedure are as follows.

A DeepLabCut network was trained on a CPU with a MacOS operating system primarily using the graphical user interface (GUI) built into DeepLabCut. ResNet 50, a pre-trained deep, residual neural network with 50 deconvolutional layers, was used as the backbone for the DeepLabCut network (Mathis et al., 2018; Nath et al., 2019). The DeepLabCut network was first trained to label videos of adult fish (wild adult fish and F1 fish at the adult timepoint) in three stages, with an additional refinement step necessary to accurately label F1 fish at the juvenile timepoint. First, 20 frames were extracted from 25 videos featuring 15 different individuals using the *extract frames* function with kmeans clustering on visual appearance to provide a diverse sample of the escape response behaviour. Additional frames were manually extracted from two of the videos to cover the entire duration of the response. The three points of interest were manually labeled on all extracted frames using the *label frames* function in the interactive GUI in DeepLabCut. Using the *check labeled frames* function, frames were exported with labels and visually inspected to ensure all labels were plotted and saved accurately. The *create training dataset* function was used to divide labeled frames into a training set (95% of frames and a test set (5% of frames). The network was trained for 300,000 iterations on the training set. Despite low mean average Euclidean error (MAE) between manual labels and labels plotted by DeepLabCut in the training set (1.72 pixels) as well as in the test set not used in the training process (1.91 pixels), the network did not generalize well upon visual inspection.

The network was refined using an expanded training set. New videos run through the network using the *analyze videos* function, where the fish did not track well, were identified by visual inspection. Using the *extract outlier frames* function with the *jump* parameter, 1-20 outlier frames from each of 48 new videos were added to the training dataset. Labels on these frames were manually adjusted using the interactive GUI in DeepLabCut. The network was trained for an additional 380,000 interactions on a training data set that now contained frames from 73 videos featuring 43 different individuals. After this second round of training, MAE in the training set (1.83 pixels) and the test set (2.35 pixels) remained low. The network showed definite improvement in generalization but still struggled to accurately label new videos.

The network was refined a second time by similarly selecting videos where the network did not track well and using the *extract outlier frames* function with the *jump* parameter to extract 1-20 outlier frames from 30 new videos and then using the DeepLabCut GUI to manually adjust labels. The network was then trained for an additional 320,000 interactions on labeled frames from 103 videos featuring 59 different individuals. Evaluation of the network resulted in a MAE of 1.82 pixels for the training set and 1.98 pixels for the test set. The network generalized well across adult fish. All videos from wild fish and F1 fish at the adult timepoint were run through this network using the *analyze videos* function with *save as CSV*, *plot trajectories*, and *create labeled videos* options set to yes.

A final refinement step was necessary to create a network that accurately labeled videos of F1 fish at the juvenile timepoint, due to the difference in size between the juvenile fish and the adult fish. Videos of F1 juvenile fish were run through the adult network and videos with poor tracking were identified by visual inspection. The *extract outlier frames* function with the *jump* parameter was used to extract 1-20 outlier frames from F1 juvenile videos and labels were manually adjusted using the DeepLabCut GUI. The network was further trained for 380,000 iterations using the additional F1 juvenile videos along with the 103 videos of adult sized fish previously used in training. The training set had an MAE of 1.62 pixels and the test set had an MAE of 1.72 pixels. The network generalized well across juvenile sized fish. All videos from the F1 juvenile timepoint were run through this network using the *analyze videos* function with *save as CSV*, *plot trajectories*, and *create labeled videos* options set to yes.
